# Brg1 regulates fibroblast-myofibroblast transition to promote renal fibrosis

**DOI:** 10.1101/2023.12.22.572996

**Authors:** Xiaoyan Wu, Yajun Luo, Aoqi Kang, Jiayao Ni, Ming Kong, Tao Zhang

## Abstract

Excessive fibrogenesis serves to disrupt the anatomical and functional integrity of the kidneys contributing to renal failure. Renal fibroblast is the major precursor to myofibroblast, the effector cell type of renal fibrosis. How fibroblast-myofibroblast transition (FMyT) is regulated in the kidneys remains incompletely understood. In the present study we investigated the role of Brahma related gene 1 (Brg1), a chromatin remodeling protein, in renal fibrosis focusing on mechanistic insights and translational potential. We report that Brg1 was up-regulated during FMyT both *in vitro* and *in vivo*. Brg1 deletion in fibroblasts partially blocked TGF-β induced FMyT *in vitro* and attenuated renal fibrosis in three different animal models. Importantly, conditional Brg1 knockout in *Postn*^+^ mature myofibroblasts mitigated renal fibrosis induced by unilateral ureteral obstruction (UUO) or ischemia-reperfusion (IR) in mice. Transcriptomic analysis uncovered Prune2 as a potential target for Brg1. Brg1 interacted with E2F1 to activate Prune2 transcription during FMyT. Concordantly, Prune2 knockdown suppressed TGF-β induced FMyT *in vitro* and dampened renal fibrosis in mice. Mechanistically, Prune2 likely contributed to FMyT by augmenting phosphorylation and activity of the pro-fibrogenic transcription factor PU.1. Finally, small-molecule Brg1 inhibitor PFI-3 exhibited strong antifibrotic potency in established models of renal fibrosis. In conclusion, our data provide compelling evidence that BRG1 is a pivotal regulator of as well as a promising therapeutic target for renal fibrosis.

## Introduction

Chronic kidney disease (CKD) is defined as the progressive loss of renal functions due to a host of etiological factors including glomerular nephropathy, diabetic nephropathy, and hypertension. Owing to the global pandemic of metabolic diseases and population aging, newly diagnosed cases of CKD have been steadily increasing in the past decade affecting more than 700 million people worldwide(1). Without effective intervention, CKD patients will eventually develop end-stage renal disease (ESRD) necessitating life-long dialysis and/or kidney transplantation(2). In the US, mortality rates in individuals 66 years or older with CKD are twice as high as those without creating significant socioeconomic burdens(3). Regardless of etiology, renal fibrosis, defined as extensive deposition of extracellular matrix proteins in the glomerulus and interstitial space, is a common pathophysiological process in CKD. Renal fibrosis leads to rigidification of renal tissues, disruption of normal renal architecture, and interference with key renal functions. Renal dampens prognosis in patients with ESRD and is considered an endpoint against which novel pharmacotherapies are devised and evaluated(4).

Myofibroblast, believed to be the effector of fibrogenesis in all organs including the kidneys, is a specialized cell type possessing both the ability to produce ECM proteins like a fibroblast cell and the ability to contract like a muscle cell(5). Myofibroblasts are absent from the kidneys under physiological conditions, only emerge during kidney injuries, and recede during the resolution of renal fibrosis(6). The origin from which myofibroblasts arise *in vivo* has been a topic of great intrigue and controversy. It was once thought that tubular epithelial cells, through a process known as epithelial-mesenchymal transition (EMT), and vascular endothelial cells, through a similar process known as endothelial-mesenchymal transition (EndMT), could trans-differentiate into myofibroblasts and thus contribute to renal fibrosis(7, 8). The development of genetic lineage tracing techniques has allowed fine mapping of the source(s) from which myofibroblasts are derived. Using a set of sophisticatedly engineered mouse strains, each of which carries a lineage-specific reporter, LeBleu et al have demonstrated that resident fibroblasts account for a majority of ECM-producing myofibroblasts (>50%) in the fibrotic kidneys(9). On the contrary, the contribution of either epithelial cells or endothelial cells to the pool of myofibroblasts appears to be negligible. More recently, with the help of a combination of genetic tracing and single-cell transcriptomic tools, Kuppe et al have shown that PDGFRa^+^/PDGFRb^+^ mesenchymal cells including all fibroblasts represent the major myofibroblast population in human kidney fibrosis(10). *In vitro* cultured fibroblasts can be programmed to differentiate into myofibroblasts by a host of stimuli including TGF-β, PDGF, Ang II, and high glucose(11).

A profound transcriptomic change is recorded during FMyT(12). In mammalian cells, transcription is acutely influenced by the epigenetic machinery, which encompasses DNA/histone modifying enzymes, non-coding RNAs, and chromatin remodeling proteins(13). Brahma related gene 1 (Brg1) is the core component of the SWI/SNF chromatin remodeling complex providing enzymatic activity for mobilization of nucleasomes(14). It has long been documented that BRG1 plays an essential role in TGF-β dependent transcription functioning as a co-factor for SMADs(15). Transcriptional events mediated by TGF-β signaling through SMAD family transcription factors are considered a paradigm in programming fibroblast-myfibroblast transition. A previous study by Gong *et al* claimed, based on immunohistochemical staining of renal tissues, that Brg1 was predominantly expressed in tubular epithelial cells(16). However, recent single-cell transcriptomic studies performed in humans and in mice indicate that Brg1 is expressed at equivalent levels in different renal cell linages(17, 18). Fibroblasts represent the predominant source of ECM-producing myofibroblasts during renal fibrosis, in which cellular compartment the role of Brg1 remains unclear. Therefore, we hypothesized that Brg1 might be involved in FMyT and renal fibrosis. Our data demonstrate that Brg1 plays an essential role in renal fibrosis by promoting fibroblast-myofibroblast transition. Prune2 is a likely downstream target of Brg1, which contributes to FMyT and renal fibrosis by activating PU.1. Finally, pharmaceutical inhibition of Brg1 with a small-molecule compound PFI-3 achieves potent antifibrotic effects in mice.

## Methods

### Animals

All animal experiments were reviewed and approved by the Ethics Committee on Humane Treatment of Laboratory Animals of Nanjing Medical University and were performed in accordance with the ethical standards laid down in the 1964 Declaration of Helsinki and its later amendments. *Smarca4*^f/f^ mice(19), *Col1a2*-Cre^ERT2^ mice(20), and *Postn*-Cre^ERT2^ mice(21) have been described previously. Renal fibrosis was induced by one of the following methods as previously described. Briefly, for the UUO procedure (22), 8-10wk male mice (25-27g) were anesthetized with isoflurane and an incision was created on the left flank beneath the ribcage to expose the left ureter. Double ligature of the ureter was performed with 5-0 Mersilk; for the sham procedure, the ureter was located and placed back without ligation. The wound was then closed using a subcuticular suture and the mice were allowed to recover on a heating pad before being returned to the cage. The mice were sacrificed 2wk after the surgery. For the ischemia-reperfusion procedure (23), 8-10wk male mice (25-27g) were anesthetized and a midline laparotomy incision was created with a scalpel. Ischemia was induced by applying a micro serrafine clip onto the left renal artery and vein for 30min. The clamp was removed and the skin was closed with silk suture. The mice were allowed to recover on a heating pad before being transferred back to the cage. For both the UUO procedure and the IR procedure, iodine/alcohol solution was applied to the surgical area to minimize infection.

To induce Cre expression in the *Col1a2*-Cre^ERT2^ mice, tamoxifen (Cat#: T2859, Sigma) was injected peritoneally (50mg/kg) for 7 consecutive days followed by a washing phase of 7 days. To induce Cre expression in the *Postn*-Cre^ERT2^ mice, tamoxifen was injected peritoneally (50mg/kg) for 5 consecutive days followed by maintaining the mice on a TAX-containing diet (Cat#: TD.130855, Inotiv) until the day when the mice were sacrificed. In certain experiments, the mice were injected peritoneally PFI-3 (Cat#: S7315, Selleck) at a dose of 50mg/kg. In certain experiments, recombinant adeno-associated virus AAV-Anc80 (24, 25) carrying Prune2 shRNA (GUGGGAUAGUCAUAUAAGUTT) downstream of the *Postn* promoter(26). The AAV was injected into C57/B6 mice intravenously (i.v.) at a dose of 1× 10^11^ vg.

### Cell culture, plasmids, and transient transfection

Primary murine renal fibroblasts were isolated and cultured as previously described(27). Primary renal tubular epithelial cells and podocytes were isolated as previously described(28, 29). *Prune2* promoter-luciferase construct was generated by amplifying genomic DNA spanning the proximal promoter and the first exon of Prune2 gene (-1500/+50) and ligating into a pGL3-basic vector (Cat#: E1751, Promega). Truncation mutants were made using a QuikChange kit (Cat#: 200518, Thermo Fisher Scientific) and verified by direct sequencing. Small interfering RNAs were purchased from Dharmacon (Cat#: LQ-016593-02-0010). Transient transfections were performed with Lipofectamine 2000 (Cat#: 11668019, Thermo Fisher Scientific). Luciferase activities were assayed 24-48 hours after transfection using a luciferase reporter assay system (Cat#: E1500, Promega) as previously described(30).

### RNA Isolation and Real-time PCR

RNA was extracted with the RNeasy RNA isolation kit (Cat#: 74104, Qiagen) as previously described(31, 32). Reverse transcriptase reactions were performed using a SuperScript First-strand Synthesis System (Cat#: 18080093, Invitrogen). Real-time PCR reactions were performed on an ABI Prism 7500 system. The primers are list in the supplementary Table I. Ct values of target genes were normalized to the Ct values of house-keeping control gene (18s, 5’-CGCGGTTCTATTTTGTTGGT-3’ and 5’-TCGTCTTCGAAACTCCGACT-3’ for both human and mouse genes) using the ΔΔCt method and expressed as relative mRNA expression levels compared to the control group which is arbitrarily set as 1.

### Protein extraction and Western blot

Whole cell lysates were obtained by re-suspending cell pellets in RIPA buffer (50 mM Tris pH7.4, 150 mM NaCl, 1% Triton X-100) with freshly added protease and phosphatase inhibitors (Cat#: 4693132001, Roche) as previously described(33). Antibodies used for Western blotting are listed in the supplementary Table II. For densitometrical quantification, densities of target proteins were normalized to those of β-actin. Data are expressed as relative protein levels compared to the control group which is arbitrarily set as 1.

### Chromatin Immunoprecipitation (ChIP)

Chromatin immunoprecipitation (ChIP) assays were performed essentially as described before(34, 35). Aliquots of lysates containing 100 μg of protein were used for each immunoprecipitation reaction with indicated antibodies followed by adsorption to protein A/G PLUS-agarose beads (Cat#: sc-2003, Santa Cruz Biotechnology). DNA-protein cross-link was reversed by heating the samples to 65 °C overnight. Proteins were digested with proteinase K (Cat#: P2308, Sigma), and DNA was phenol/chloroform-extracted and precipitated by 100% ethanol. Precipitated genomic DNA was amplified by real-time PCR. A total of 10% of the starting material is also included as the input.

### Histology

Histological analyses were performed essentially as described before(36). Picrosirius Red staining was performed with a commercially available kit (Cat#: ab150681, Abcam) per vendor recommendation. Paraffin sections were de-waxed and hydrated before being immersed in the Picrosirius Red solution for 60 minutes under room temperature. After two washes with acidified water, the slides were rinsed with 100% ethanol. The slides were further dehydrated twice with 100% ethanol before being cleared and mounted for visualization. Masson’s trichrome staining was performed with a commercially available kit (Cat#: HT15-1KT, Sigma) per vendor recommendation. Briefly, paraffin sections were de-waxed and hydrated before being mordanted in Bouin’s Solution at 56°C for 15 minutes. The slides were washed with running tap water to remove yellow color and then stained with Weigert’s Iron Hematoxylin Solution for 5 minutes. After several washes and rinses, the slides were stained with sequentially with Biebrich Scarlet-Acid Fucshin Solution, Phosphotungstic/Phosphomolybdic Acid Solution, and Aniline Blue Solution. After several more washes and rinses, the slides were dehydrated 100% ethanol before being cleared and mounted for visualization. Pictures were taken using an Olympus IX-70 microscope. Quantifications were performed with Image J. For each mouse, at least three slides were stained and at least five different fields were analyzed for each slide.

### EdU incorporation assay

Cell proliferation was evaluated by 5-ethynyl-2’-deoxyuridine (EdU) incorporation assay as previously described(37). Briefly, cells were seeded in triplicates in 48-well plates. EdU solution (Cat#: C10086, Thermo Fisher) was added to and incubated with the cells for 18h. After nuclear staining with DAPI for 30 min at room temperature, the fluorescence at 594 nm was detected with an Olympus IX-70 microscope. EdU positive cells from at least 5 randomly chosen fields were counted for each triplicate well. Data are expressed as relative EdU staining normalized to the control group arbitrarily set as 1.

### Boyden chamber trans-well assay

The cells were trypsinized and seeded into Boyden chambers (PET track-etched, 8-μm pores, 24-well format; Cat#: 353097, Becton Dickinson) in serum-free DMEM medium. Complete culture medium containing 10% FBS was added to the lower chamber. The cells migrating from the upper chamber were fixed with 4% paraformaldehyde, stained with 0.1% crystal violet, and counted under a microscope. Cell numbers from 5 random fields were counted in each well. Data are expressed as relative number of migrated cells normalized to the control group arbitrarily set as 1.

### Collagen contraction assay

The cells were trypsinized, mixed with 4x the volume of Collagen Gel Working Solution (Cat#: 354236, Corning) and incubated for 1 hr at 37°C. 4. After collagen polymerization, 1.0 mL of culture medium was added atop. The collagen gel size change was measured 24h later and quantified with Image Pro Plus. Data are expressed as relative contraction normalized to the control group arbitrarily set as 1.

### RNA Sequencing and Data Analysis

RNA-seq was performed as previously described(38). Total RNA was extracted using the TRIzol reagent according to the manufacturer’s protocol. RNA purity and quantification were evaluated using the NanoDrop 2000 spectrophotometer (Thermo Scientific, USA). RNA integrity was assessed using the Agilent 2100 Bioanalyzer (Agilent Technologies, Santa Clara, CA, USA). Then the libraries were constructed using TruSeq Stranded mRNA LT Sample Prep Kit (Illumina, San Diego, CA, USA) according to the manufacturer’s instructions and sequenced on an Illumina HiSeq X Ten platform and 150 bp paired-end reads were generated. Raw data (raw reads) of fastq format were firstly processed using Trimmomatic and the low quality reads were removed to obtain the clean reads. The clean reads were mapped to the mouse genome (Mus_musculus.GRCm38.99) using HISAT2. FPKM of each gene was calculated using Cufflinks, and the read counts of each gene were obtained by HTSeqcount. Differential expression analysis was performed using the DESeq (2012) R package. P value < 0.05 and fold change > 1.5 or fold change < 0.66 was set as the threshold for significantly differential expression. Hierarchical cluster analysis of differentially expressed genes (DEGs) was performed to demonstrate the expression pattern of genes in different groups and samples. GO enrichment and KEGG pathway enrichment analysis of DEGs were performed respectively using R based on the hypergeometric distribution.

### Statistical analysis

For comparison between two groups, two-tailed t-test was performed. For comparison among three or more groups, one-way ANOVA or two-way ANOVA with post-hoc Turkey analyses were performed using an SPSS package. The assumptions of normality were checked using Shapiro-Wilks test and equal variance was checked using Levene’s test; both were satisfied. *p* values smaller than .05 were considered statistically significant (*). All *in vitro* experiments were repeated at least three times and three replicates were estimated to provide 80% power.

## Results

### Brg1 regulates fibroblast-myofibroblast transition in vitro

When primary renal fibroblasts were exposed to TGF-β, a potent inducer of FMyT, robust induction of α-SMA (encoded by *Acta2*), a prototypical myofibroblast marker, was detected by qPCR (Fig.1A) and Western blotting (Fig.1B), indicative of FMyT taking place. Of interest, there was a concomitant increase in Brg1 expression at both mRNA (Fig.1A) and protein (Fig.1B) levels. Importantly, when primary renal fibroblasts were isolated from the mice in which renal fibrosis was developing as a result of either the unilateral ureteral obstruction (UUO) procedure or the ischemia-reperfusion (IR) procedure, expression of levels of Brg1 and α-SMA followed almost identical kinetics (Fig.S1).

**Figure 1:**
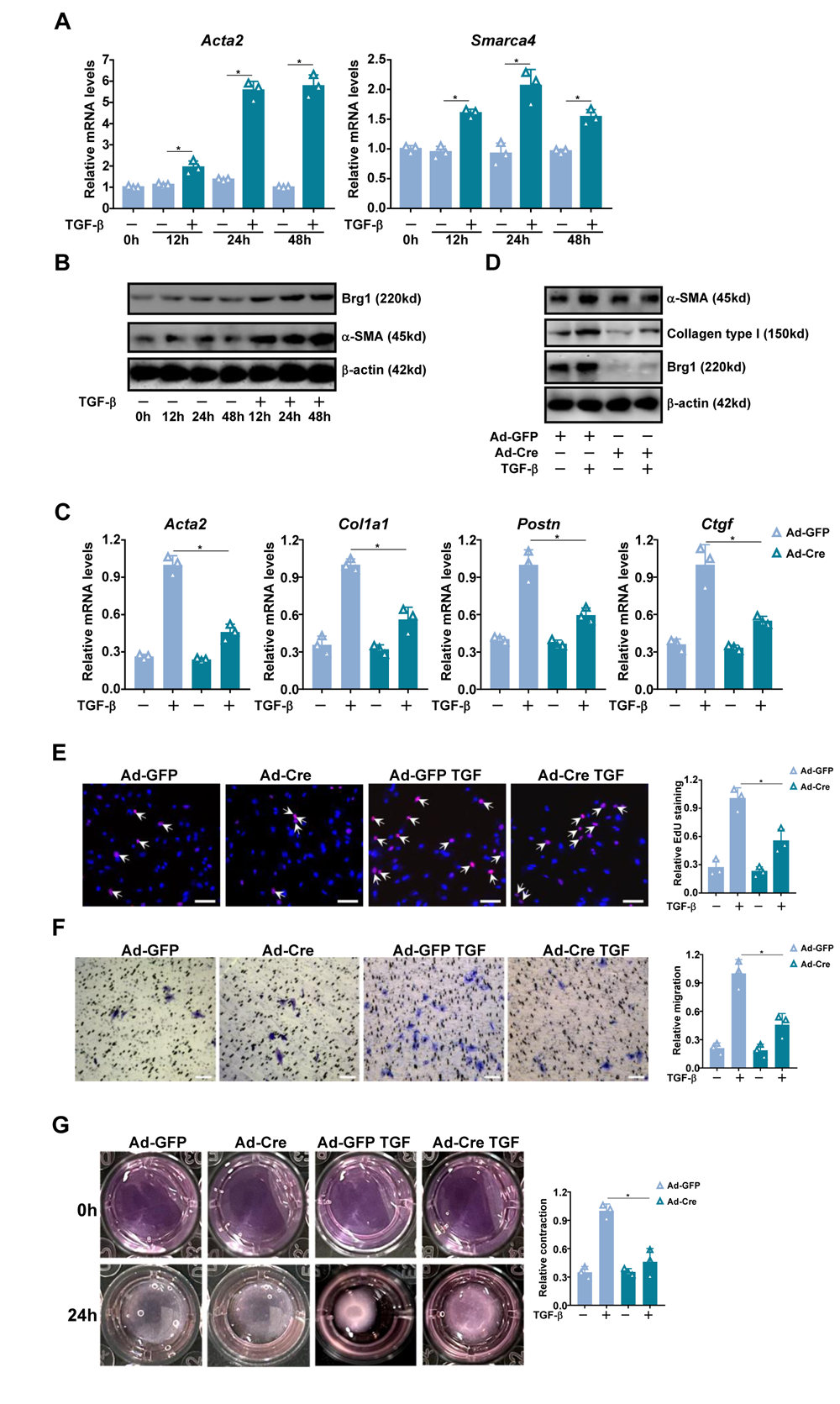
Brg1 regulates fibroblast-myofibroblast transition in vitro. (**A, B**) Primary renal fibroblasts were treated with TGF-β (2ng/ml) and harvested at indicated time points. Brg1 expression was examined by qPCR and Western blotting. (**C-G**) Primary renal fibroblasts isolated from *Brg1*^f/f^ mice were transduced with Ad-GFP or Ad-Cre followed by treatment with TGF-β. Myofibroblast marker genes were examined by qPCR (C) and Western blotting (D). Proliferation was evaluated by EdU incorporation (E). Migration was evaluated by Boyden chamber assay (F). Collagen contraction assay (G). Scale bar, 50μm

Next, primary renal fibroblasts were isolated from the Brg1-flox mice and Brg1 deletion was achieved by the addition of Cre enzyme: Brg1 deletion markedly dampened TGF-β induced FMyT as evidenced by expression levels of myofibroblast marker genes (Fig.1C, 1D) and the abilities to proliferate (Fig.1E), migrate (Fig.1F), and contract (Fig.1G). On the contrary, over-expression of Brg1 strongly enhanced FMyT (Fig.S2). Taken together, these data suggest that Brg1 might play an essential role in FMyT *in vitro*.

### Fibroblast-specific Brg1 ablation attenuates renal fibrosis in mice

To delete Brg1 in resident fibroblasts, *Brg1*^f/f^ mice were crossed to the *Col1a2*-Cre^ERT2^ mice. The Brg1^FCKO^ mice (*Brg1*^f/f^; *Col1a2*-Cre^ERT2^) and WT mice (*Brg1*^f/f^) were injected with tamoxifen to allow Cre-mediated homologous recombination (Fig.2A). Brg1 expression was significantly down-regulated in the fibroblast fraction, but not the non-fibroblast fraction, in the Brg1^FCKO^ mice compared to the WT mice (Fig.S3). These mice were then subjected to the UUO procedure to induce renal fibrosis. Brg1 deficiency in fibroblasts did not alter renal filtration function because no difference in plasma BUN levels (Fig.2B) and creatinine levels (Fig.2C). However, picrosirius red staining and Masson’s staining showed a decrease in extracellular matrix proteins in the kidneys of the Brg1^FCKO^ mice compared to the WT mice (Fig.2D). QPCR (Fig.2E) and Western blotting (Fig.2F) confirmed that Brg1 deletion in fibroblasts diminished pro-fibrogenic/myofibroblast marker genes. In addition, hydroxyproline quantification pointed to a decrease in collagenous tissue deposition (Fig.2G). The conclusion that fibroblast-specific Brg1 contributes to renal fibrosis was further validated in an alternative model of renal fibrosis induced by kidney ischemia-reperfusion (Fig.S4).

**Figure 2:**
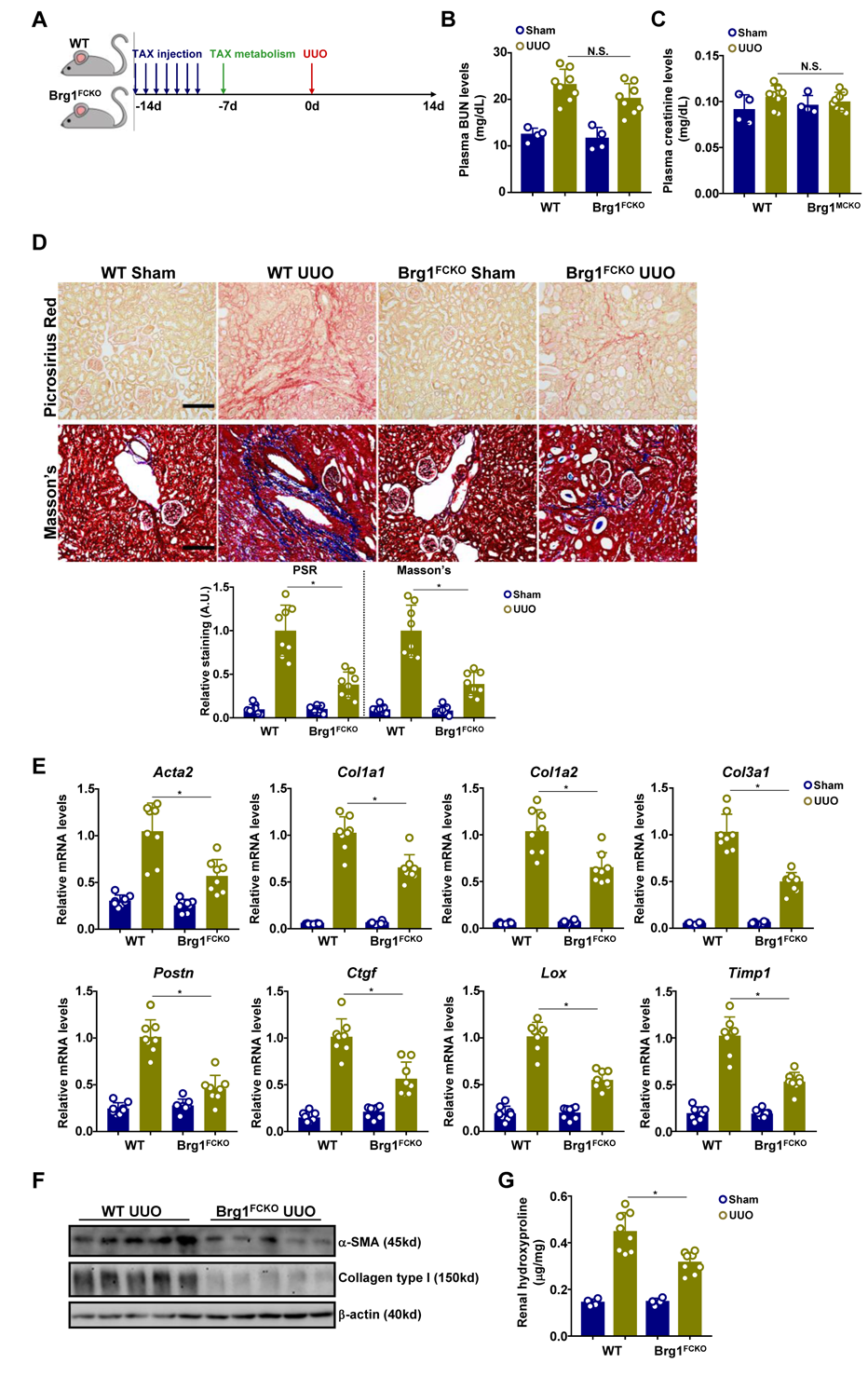
Fibroblast-specific Brg1 ablation attenuates renal fibrosis in mice. WT and Brg1^FCKO^ mice were subjected to the UUO procedure as described in Methods. (**A**) Scheme of protocol. (**B**) Plasma BUN levels. (**C**) Plasma creatinine levels. (**D**) Paraffin sections were stained with Picrosirius Red and Masson’s trichrome. (**E, F**) Pro-fibrogenic genes were examined by qPCR and Western blotting. (**G**) Hydroxyproline levels. N=8 mice for each group. Data are expressed as the means ± SD. *, *p*>0.05.

### Myofibroblast-specific Brg1 ablation attenuates renal fibrosis in mice

Because the ability of Brg1 to regulate renal fibrosis might not entirely reside in fibroblasts, the next set of experiments was performed to determine whether Brg1 deficiency in mature myofibroblast would similarly attenaute renal fibrosis. To this end, *Brg1*^f/f^ mice were crossed to the *Postn*-Cre^ERT2^ mice to generate myofibroblast conditional Brg1 knockout mice (Brg1^MCKO^, Fig.3A). Following the UUO surgery, neither plasma BUN levels (Fig.3B) nor plasma creatinine levels (Fig.3C) were significantly altered by Brg1 deletion in myofibroblasts. However, there were clearly fewer extracellular matrix proteins accumulated in the kidneys of the Brg1^MCKO^ mice than in the WT mice as evidenced by picrosirius red staining and Masson’s staining (Fig.3D). Measurements of pro-fibrogenic/myofibroblast markers by qPCR (Fig.3E) and Western blotting (Fig.3F) and quantification of hydroxyproline levels (Fig.3G) provided additional support for the notion that Brg1 might contribute to renal fibrosis by maintaining the myofibroblast phenotype. Additionally, this notion was further authenticated in the kidney ischemia-reperfusion model (Fig.S5).

**Figure 3:**
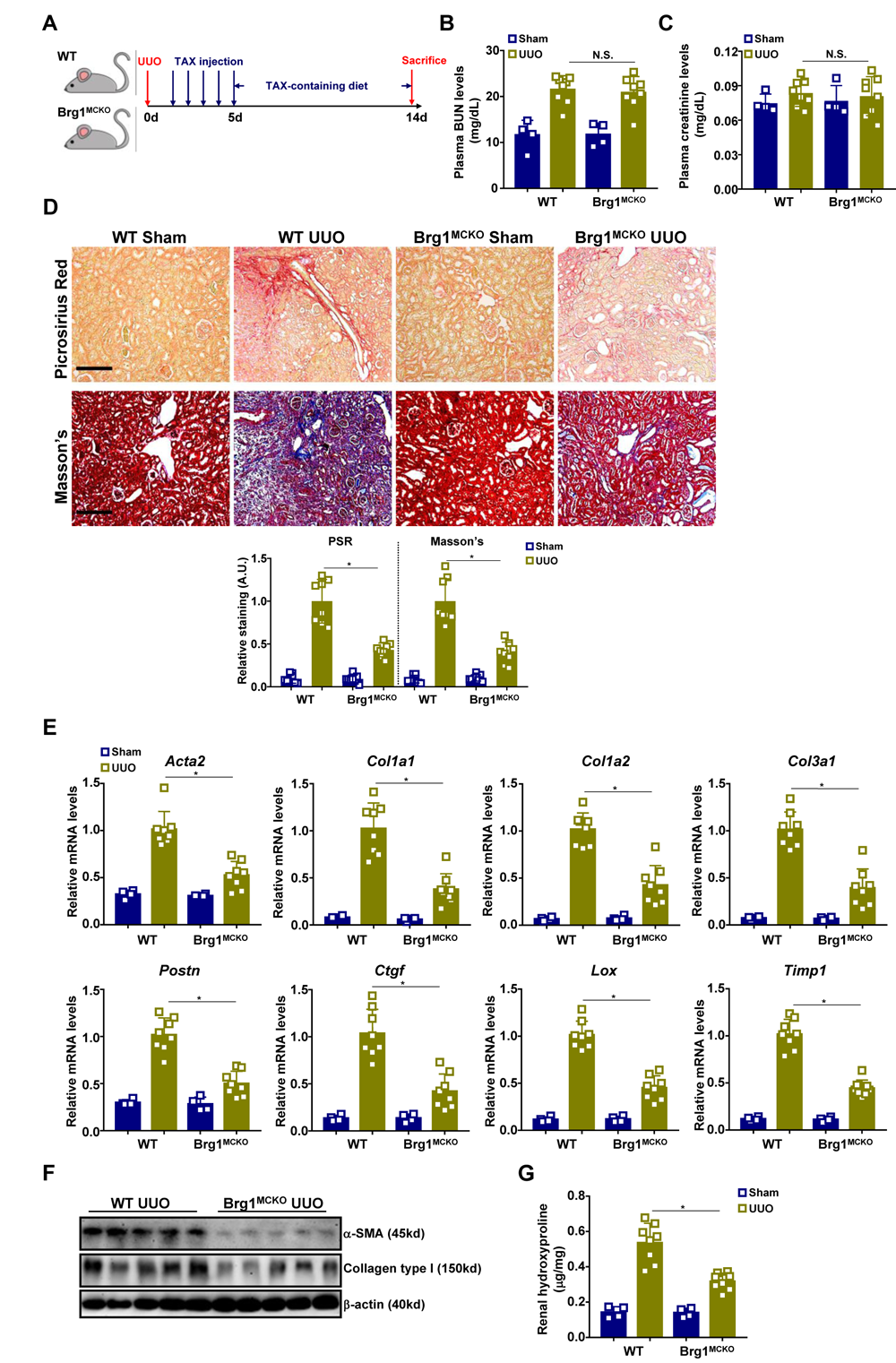
Myofibroblast-specific Brg1 ablation attenuates renal fibrosis in mice. WT and Brg1^MCKO^ mice were subjected to the UUO procedure as described in Methods. (**A**) Scheme of protocol. (**B**) Plasma BUN levels. (**C**) Plasma creatinine levels. (**D**) Paraffin sections were stained with Picrosirius Red and Masson’s trichrome. (**E, F**) Pro-fibrogenic genes were examined by qPCR and Western blotting. (**G**) Hydroxyproline levels. N=4-8 mice for each group. Data are expressed as the means ± SD. *, *p*>0.05.

### Brg1 regulates fibroblast transcriptome during FMyT

In order to identify transcription target(s) for Brg1 during FMyT, RNA-seq was performed to compare the transcriptome of wild type (Ad-GFP) and Brg1-null (Ad-Cre) renal fibroblasts (Fig.4A). Brg1 deficiency resulted in far more genes being down-regulated (909) than being up-regulated (123) using 1.5x fold change as a cut-off consistent with the notion that Brg1 primarily acts as an activator of transcription (Fig.4B). GO and KEGG analyses indicated that most genes affected by Brg1 deficiency were involved in the regulation of cell proliferation, migration, contraction, and ECM production (Fig.4C). Prune2 (also known as Bmcc1) was among the top 5 genes most significantly down-regulated by Brg1 deletion (Fig.4D). Because Prune2 has previously been detected as one of top up-regulated genes by both TGF-β and CTGF in renal fibroblasts(39), we focused on Prune2 for the remainder of the study.

**Figure 4:**
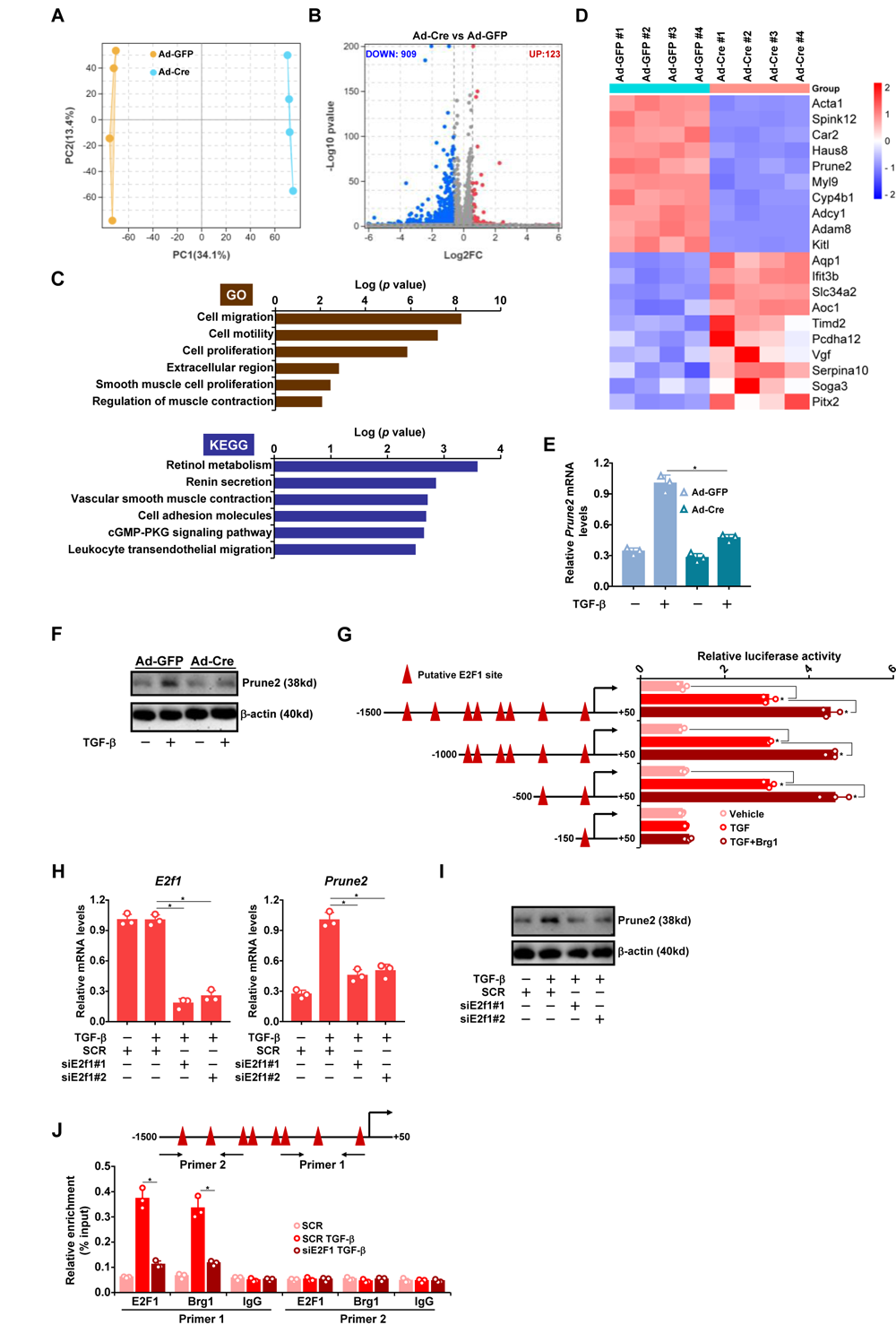
Brg1 regulates fibroblast transcriptome during FMyT. (**A-D**) Primary renal fibroblasts isolated from *Brg1*^f/f^ mice were transduced with Ad-GFP or Ad-Cre followed by treatment with TGF-β. RNA-seq was performed as described in Methods. (A) PCA plot. (B) Volcano plot. (C) GO and KEGG analyses. (D) Heatmap of top differentially expressed genes. (**E, F**) Primary renal fibroblasts isolated from *Brg1*^f/f^ mice were transduced with Ad-GFP or Ad-Cre followed by treatment with TGF-β. Prune2 expression was examined by qPCR and Western blotting. (**G**) Prune2 promoter-luciferase constructs were transfected into mouse embryonic fibroblasts (MEFs) with or without Brg1 followed by treatment with TGF-β. Luciferase constructs were normalized by protein concentration and GFP fluorescence. (**H-J**) Primary murine renal fibroblasts were transfected with indicated siRNAs followed by treatment with TGF-β. Prune2 expression in the kidney tissues was examined by qPCR and Western blotting. ChIP assays were performed with anti-E2F1, anti-Brg1, or IgG.

Prune2 expression was up-regulated during FMyT *in vivo* in two different models of renal fibrosis (Fig.S6A, S6B). Prune2 expression was also inducible with TGF-β treatment (Fig.S6C). Brg1 deletion dampened (Fig.4E, 4F) whereas Brg1 over-expression enhanced (Fig.S7) Prune2 induction by TGF-β. Further, Prune2 levels were found to be lower in the kidneys of the Brg1^MCKO^ mice than in the WT mice (Fig.S8). A series of E2F1 sites were identified within the Prune2 promoter region (Fig.4G). Reporter assay showed that TGF-β treatment augmented the activity of the Prune2 promoter, which was further enhanced by Brg1 over-expression. However, when progressive deletions went beyond -150bp relative to the transcription start site (TSS), the Prune2 promoter failed to respond to either TGF-β treatment or Brg1 over-expression (Fig.4G) suggesting that E2F1 motif located between -500 and -150 might be responsible for recruiting Brg1 to mediate Prune2 induction in response to TGF-β. E2F1 knockdown indeed abrogated Prune2 induction by TGF-β (Fig.4H, 4I). ChIP assay confirmed that both Brg1 and E2F1 were recruited to the Prune2 promoter and that E2F1 knockdown dampened the occupancies of both E2F1 and Brg1 (Fig.4J).

### Prune2 is essential for fibroblast-myofibroblast transition in vitro and renal fibrosis in vivo

In order to determine the role of Prune2 might play in FMyT and renal fibrosis, the following experiments were performed. Prune2 knockdown by two different pairs of siRNAs down-regulated the expression of myofibroblast marker genes (Fig.5A), decreased proliferation (Fig.5B), diminished migration (Fig.5C), and attenuated contraction (Fig.5D) of renal fibroblasts exposed to TGF-β. It was observed that Prune2 silencing did not appear to influence TGF-β signaling as measured by SMAD3 phosphorylation (Fig.S9). Next, shRNA targeting Prune2 (shPrune2) was placed downstream of the *Postn* promoter and packaged into AAV-Anc80. C57/B6 mice were injected with AAV carrying shPrune2 or a control vector (shC) followed by the UUO procedure to induce renal fibrosis (Fig.5E). QPCR measurements confirmed that Prune2 expression was specifically down-regulated in the fibroblasts, but not in tubular epithelial cells or podocytes, by AAV-shPrune2 in the kidneys (Fig.S10). Myofibroblast-specific Prune2 knockdown in mice did not appreciably alter renal filtration function as evidenced by comparable levels of plasma BUN (Fig.5F) and plasma creatinine (Fig.5G). However, renal fibrosis was markedly ameliorated in the shPrune2 mice compared to the shC mice as evidenced by histological staining (Fig.5H), by qPCR (Fig.5I) measurements of myofibroblast marker genes, and by hydroxyproline quantification (Fig.5J). Likewise, Prune2 deletion in the myofibroblasts mollified renal fibrosis in an alternative model induced by ischemia-reperfusion (Fig.S11).

**Figure 5:**
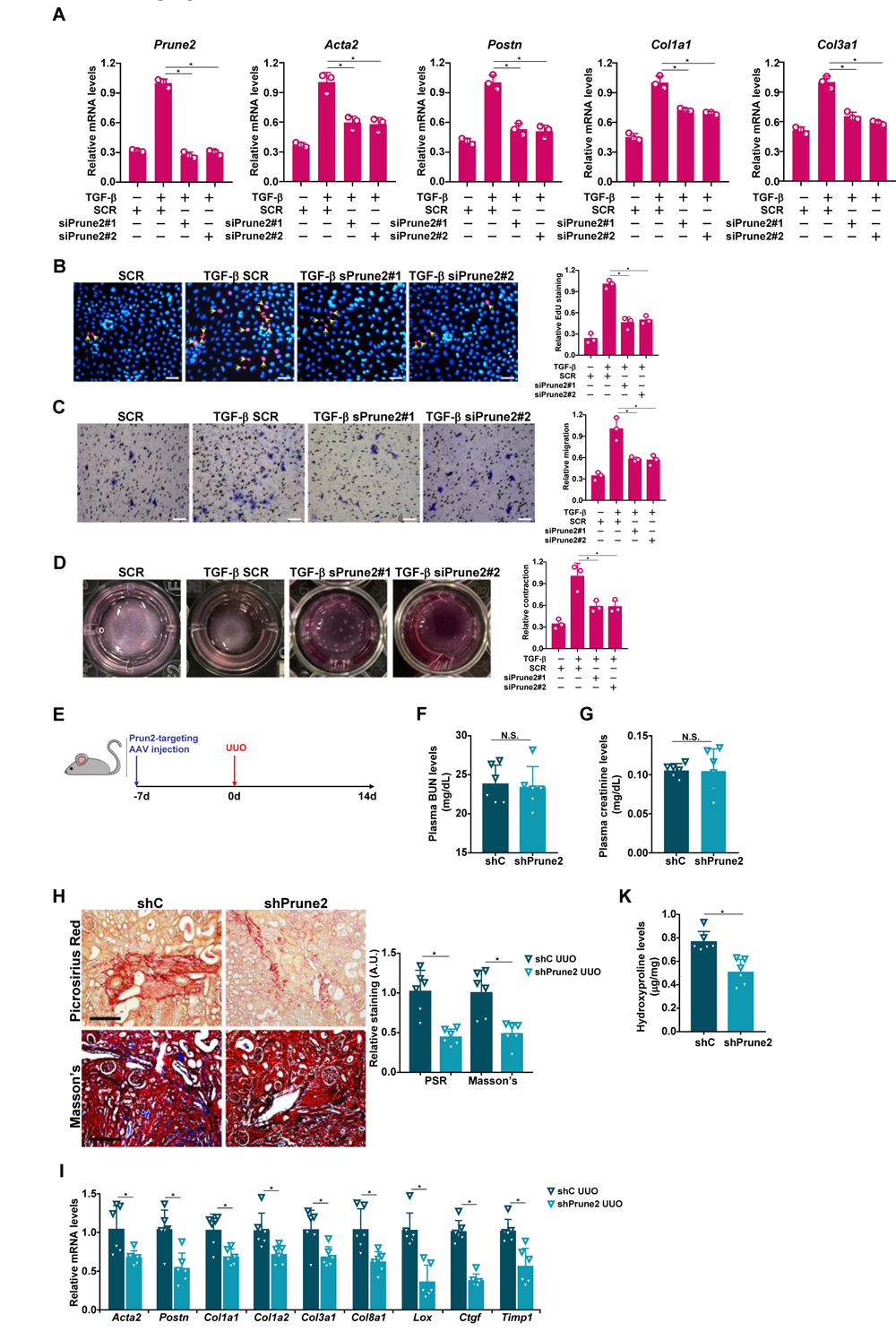
Prune2 is essential for fibroblast-myofibroblast transition in vitro and renal fibrosis in vivo. (**A-D**) Primary renal fibroblasts were transfected with siRNAs targeting Prune2 or scrambled siRNAs (SCR) followed by treatment with TGF-β. Myofibroblast marker genes were examined by qPCR (A). Proliferation was evaluated by EdU incorporation (B). Migration was evaluated by Boyden chamber assay (C). Collagen contraction assay (D). Scale bar, 50μm (**E-K**) C57/BL6 mice were injected with AAV-Anc80 carrying shRNA targeting Prune2 (shPrune2) or control shRNA (shC) followed by the UUO procedure to induce renal fibrosis. Scheme of protocol (E). Plasma BUN levels (F). Plasma creatinine levels (G). Paraffin sections were stained with Picrosirius Red and Masson’s trichrome (H). Pro-fibrogenic genes were examined by qPCR (I). Hydroxyproline levels (J). N=6 mice for each group. Data are expressed as the means ± SD. *, *p*>0.05.

### Prune2 regulates FMyT through activating PU.1

RNA-seq was performed to gain a genomewide perspective on the mechanism whereby Prune2 might contribute to FMyT. Prune2 knockdown significantly altered the transcriptome of renal fibroblasts (Fig.6A). In all, 602 genes met the threshold (1.5x fold change and *p*<0.05), of which 443 were down-regulated and 159 were up-regulated (Fig.6B). GO analysis indicated that Prune2 influenced the expression of genes that might regulate cell proliferation, migration and ECM production (Fig.6C). Among the top ranked genes altered by Prune2 knockdown there were prototypical myofibroblast marker genes (e.g., *Acta2*, *Postn*, and *Col8a1*), ECM remodeling enzymes (e.g., *Mmp9*, *Mmp12*, *Mmp13*, and *Ctss*), redox enzymes (e.g., *Ncf1*, *Nqo1*, and *Gsta3*), and some of the well-established antifibrotic transcription factors (e.g., *Irf9*, *Stat1*, and *Stat2*) (Fig.6D). Of interest, HOMER analysis revealed that Prune2 deficiency most significantly dampened the activity of PU.1 (Fig.6E), a transcription factor recently identified as the master regulator of tissue fibrosis(40).

**Figure 6:**
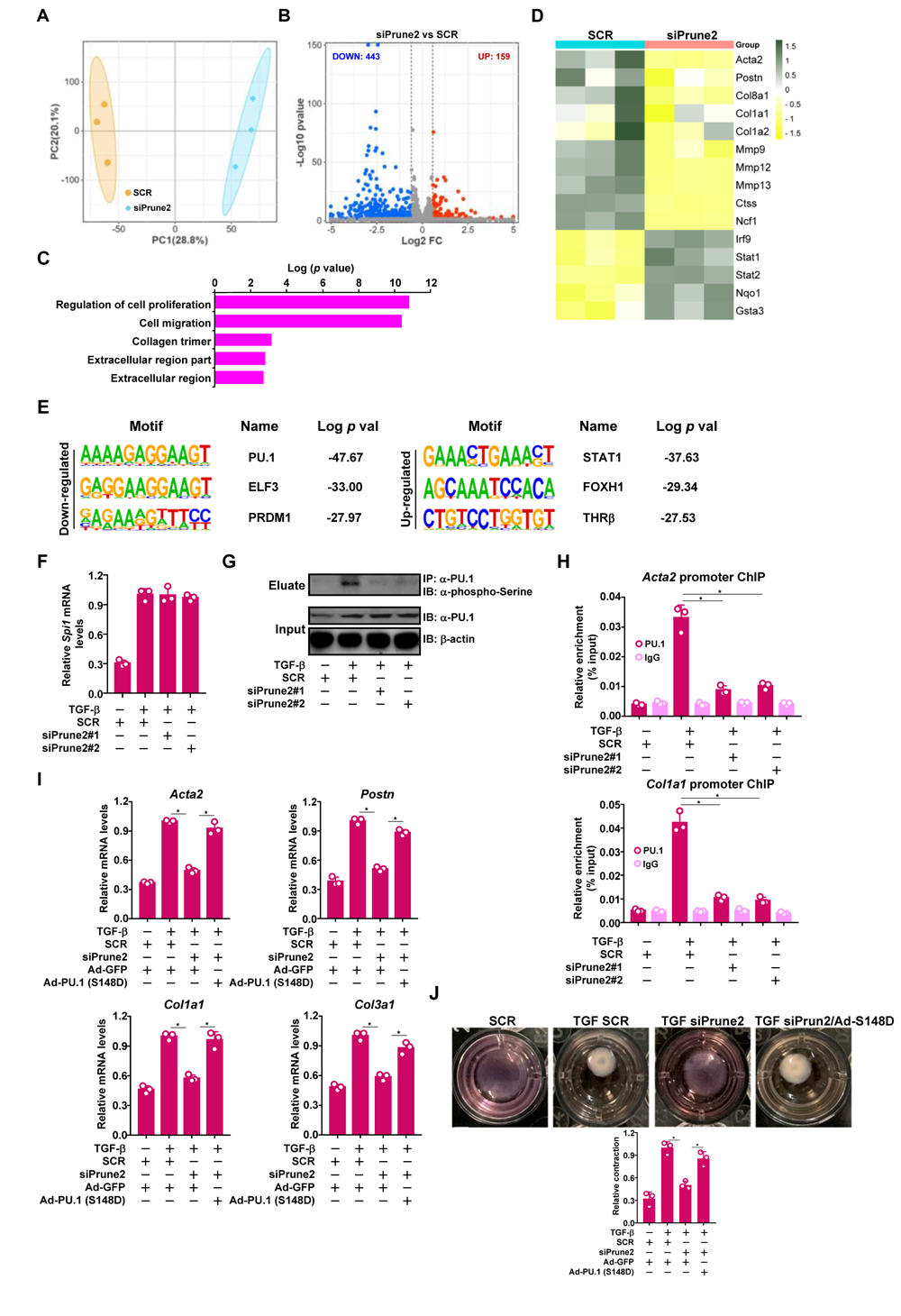
Prune2 regulates FMyT through activating PU.1. (**A-E**) RNA-seq was performed as described in Methods. (A) PCA plot. (B) Volcano plot. (C) GO analyses. (D) Heatmap of top differentially expressed genes. (E) HOMER analysis. (**F, G**) Primary renal fibroblasts were transfected with siRNAs targeting Prune2 or scrambled siRNAs (SCR) followed by treatment with TGF-β. PU.1 expression was examined by qPCR. Whole cell lysates were immunoprecipitated with anti-PU.1.

PU.1 (encoded by *Spi1*) expression levels were up-regulated by TGF-β treatment but remain unaffected by Prune2 knockdown (Fig.6F). However, PU.1 phosphorylation was sensitive to both TGF-β treatment and Prune2 knockdown (Fig.6G). Consistently, PU.1 binding to the *Acta2* promoter and the *Col1a1* promoter was dampened by Prune2 knockdown (Fig.6H). Phosphorylation of PU.1 by casein kinase 2 (CK2) at serine 148 has been shown to be required for its transcriptional activity(41–43). In order to further validate the model in which Prune2 might contribute to FMyT by modulating PU.1 phosphorylation, renal fibroblasts depleted of Prune2 were reconstituted with a mutant PU.1 in which S148 was replaced by an aspartic acid (D). As shown in Fig.6I and 6J, reconstitution of the PU.1 phospho-mimetic rescued the deficiency of FMyT caused by Prune2 knockdown.

### Pharmaceutical inhibition of Brg1 attenuates renal fibrosis in mice

We finally explored the translational potential of our finding by testing the efficacy of a small-molecule Brg1 inhibitor PFI-3 in established models of renal fibrosis (Fig.7A). Administration of PFI-3 following the UUO procedure significantly improved renal function as evidenced by reduced levels of plasma BUN (Fig.7B) and plasma creatinine (Fig.7C). In addition, fibrogenesis in the kidneys was significantly suppressed by PFI-3 as measured by picrosirius red/Masson’s staining of ECM proteins (Fig.7D), by expression levels of myofibroblast marker genes (Fig.7E, 7F), and by quantification of collagenous tissues (Fig.7G). The efficacy of PFI-3 as a potential antifibrotic reagent was further authenticated in an additional model of renal fibrosis (Fig.S12). Consistently, PFI-3 treatment antagonized TGF-β induced FMyT in cultured renal fibroblasts (Fig.S13). RNA-seq analysis confirmed that Brg1 inhibition by PFI-3 led to changes in genes involved in FMyT (Fig.S14).

**Figure 7:**
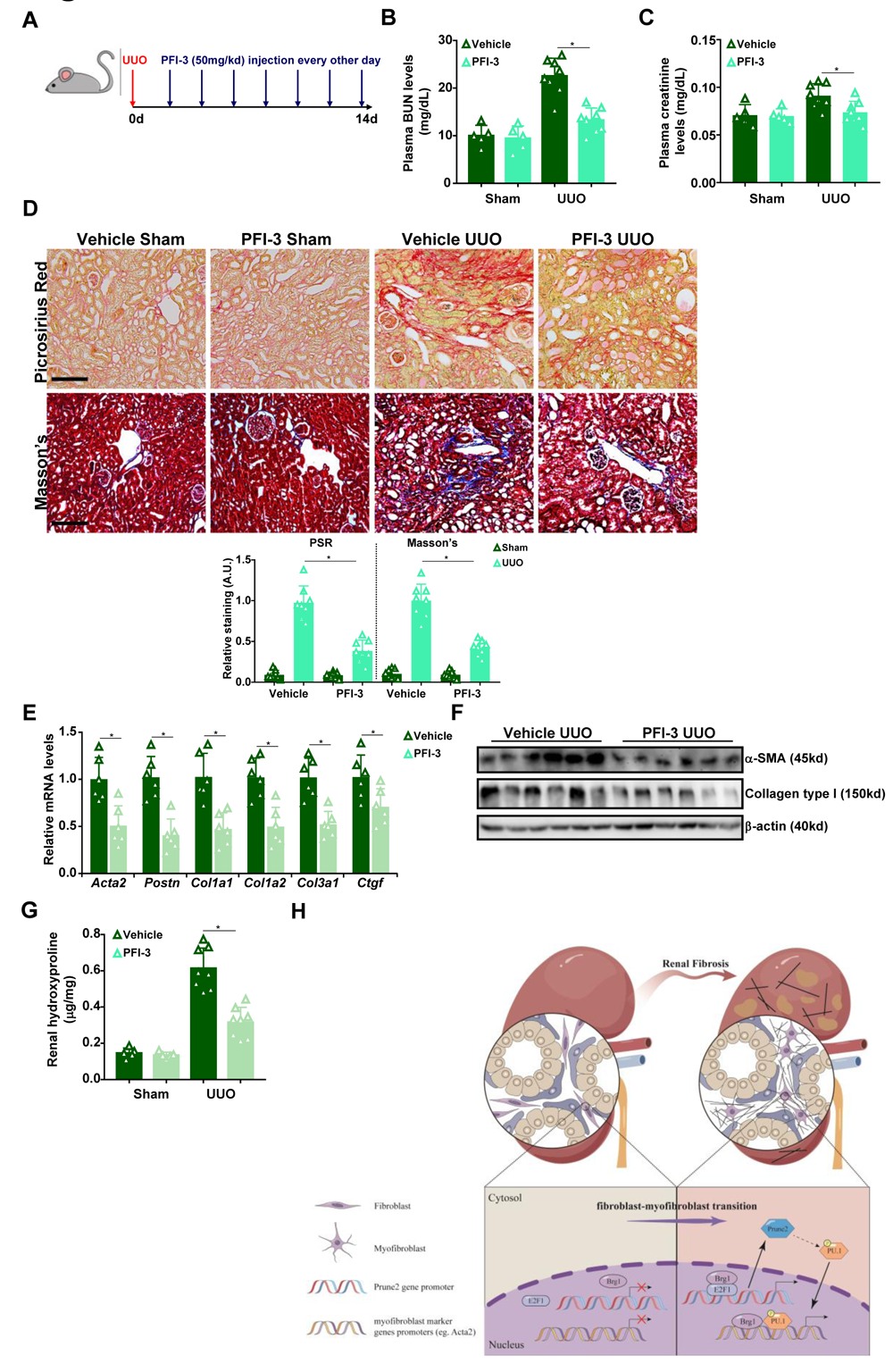
Pharmaceutical inhibition of Brg1 attenuates renal fibrosis in mice. (A-G) C57/BL6 mice were subjected to the UUO procedure to induce renal fibrosis. PFI-3 was given via peritoneal injection every other day following the surgery. Scheme of protocol (A). Plasma BUN levels (B). Plasma creatinine levels (C). Paraffin sections were stained with Picrosirius Red and Masson’s trichrome (D). Pro-fibrogenic genes were examined by qPCR (E) and Western blotting (F). Hydroxyproline levels (G). N=5-8 mice for each group. Data are expressed as the means ± SD. *, *p*>0.05. (**H**) A schematic model summarizing the key findings.

## Discussion

Renal fibroblasts represent the major cell lineage from which myofibroblasts in the fibrotic kidneys are derived through a process known as fibroblast-myofibroblast transition (FMyT). Here we detail a novel pathway in which the chromatin remodeling protein Brg1 regulates renal fibrosis by promoting FMyT. Brg1 has a relatively well-documented role in the regulation of contractile proteins in smooth muscle cells. For instance, Zhou *et al* have previously reported that forced expression of myocardin/myocardin-related transcription factors, master regulators of the contractile phenotype, in non-muscle cells leads to marked up-regulation of contractile proteins (e.g., α-SMA), to which process Brg1 serves as the rate-limiting factor(44). On the other hand, no study has systemically examined the role of Brg1 in renal fibroblasts up to this point although Brg1 has shown to interact with multiple transcription factors/co-factors involved in ECM production including SMAD(15), SRF(45), β-catenin/TCF4(46), and TEAD(47). We show here that deletion of Brg1 in either fibroblasts or myofibroblasts attenuates renal fibrosis in mice without significantly altering renal function. This is conspicuously different from previous reports showing that endothelial-specfic Brg1 deletion assuages both renal fibrosis and renal injury(22, 48). Because the contribution of endothelial cell lineage to the pool of myofibroblasts is considered marginal, these observations collectively suggest that myofibroblasts, at least those originiating from fibroblasts, play a dedicated role in producting ECM. Another implicit message taken from these observations is that Brg1 may play lineage-seletive and distinguishable roles in the pathogenesis of renal diseases. Since postnatal Brg1 deletion in mice appears to be compatible with normal life activities, it is safe to assume that indiscriminate Brg1 inhibition would engender multiple desirable effects.

Our data demonstrate that Prune2, a downstream target of Brg1, promotes FMyT and renal fibrosis. This observation is consistent with a previous report by Johnson *et al* that shows a positive correlation between Prune2 expression and treatment with pro-fibrogenic growther factors(39). Our data suggest that Prune2 likely regulates FMyT by enabling CK2-mediated PU.1 phosphorylation and activation. Multiple independent investigations have built a compelling case for the involvement of CK2 in cellular fibrogenic response. For instance, Zhang *et al*(49) and You *et al*(50) have reported that CK2 deletion or inhibition blunted TGF-β induced FMyT *in vitro*. In addition, CK2 knockdown in *db*/*db* mice appears to ameliorate renal fibrosis(51); whether this is achieved through dampening FMyT is not clear. The exact mechanism whereby Prune2 activates PU.1 remains murky at this point. Prune2 contains a BNIP2 and Cdc42GAP homology (BCH) domain that can mediate its interaction with guanine nucleotide exchange factors (GEFs) to modulate Rho activity(52) and Akt activity(53). Rho(54) and Akt(55) can potentially regulate CK2 activity thus providing a link between Prune2 and PU.1. Alternatively, there is also evidence to suggest that Rho(56) and Akt(57) may directly stimulate PU.1 activity. PU.1 inhibitor is available and has been tested in different models of tissue fibrosis(40, 58). It would be of interest to determine whether PU.1 inhibition could alter renal fibrosis *in vivo*. Clearly, more studies are needed to solve these lingering issues.

We focus on Prune2 as the major Brg1 target that mediates its pro-fibrogenic effects. There are other Brg1 targets that deserve to be closely examined for their roles in renal fibrosis in future studies. For instance, ADAM8 expression has been shown to respond to pro-fibrogenic stimuli in lung fibroblasts(59), cardiac fibroblasts(60), and dermal fibroblasts(61). Consistently, genetic deletion or pharmaceutical inhibition of ADAM8 attenuates fibrosis in the lungs(62) and in the skull(63). Kitl (also known as stem cell factor/SCF) is another interesting candidate that likely contributes to Brg1-dependent FMyT. There is evidence to show that Kit1, along with its cognate receptor Kit, promotes FMyT in a paracrine fashion by triggering the differentiation of neighboring fibroblasts; accordingly, Kitl neutralization attenuates bleomyocin-induced pulmonary fibrosis in mice(64). Of intrigue, circulating Kitl levels have been identified to be positively correlated with renal intestitial fibrosis but negatively correlated with renal fibrosis in healthy aging adults(65) and in patients with glomerulonephritis(66). Additional investigations are warranted to establish a causal relationship between these molecules and renal fibrosis.

The most important finding of the present study is that pharmaceutical inhibition of Brg1 by PFI-3 is associated with mollification of renal fibrosis as well as improvement of renal function. This observation again argues for lineage-specific roles of Brg1 in regulating renal pathophysiology (e.g., fibroblast/myofibroblast vs endothelial or macrophage). PFI-3 was designed to specifically target proteins with type VIII bromodomain, which include both Brg1 and Brm(67). Brm-null mice are viable and larger in size compared to wild type littermates owing to increased cell proliferation(68); little is known regarding the role of Brm in FMyT or tissue fibrosis. Contractile genes, which constitute part of the myofibroblast signature, seem to show varied dependence on Brg1 and Brm(69). More work is needed to delineate the redundancy of Brg1 and Brm in this process. To date, JQ1, the specific inhibitor targeting type V bromodomain protein BRD4, is among the most potent antifibrotic reagents in organ fibrosis(70). Our data reinforce the notion that bromodomain-containing chromatin remodeling proteins are master regulators of fibrogenesis. It is tempting to speculate that dual-targeting these proteins might achieve the holy grail to lower mortalities caused by fibrosis-associated organ failure.

Despite the advances proffered by the present study, major limitations caution future endeavors that aim to exploit Brg1 as a potential target to treat CKD. First, CKD typically develops in the course of years, if not decades, whereas the present study draws its conclusion primarily from model animals in which renal fibrosis develops in the course of weeks. Without strong evidence from population studies and/or CKD patients, it remains to be seen whether targeting Brg1 would bring out long-term protective/therapeutic effects. Second, although preliminary data suggest that Prune2 might be a novel and attractive target for the intervention of renal fibrosis the effort to validate Prune2 was not as exhaustive as Brg1. For instance, the kidneys are notoriously difficult for viral transduction. Even with the most specific/efficiency AAV targeting system (AAV-Anc80) available(25), the conclusion that fibroblast/myofibroblast-derived Prune2 is critical for renal fibrosis *in vivo* still needs further verification by genetic approaches (e.g., the Cre-LoxP system). Third, our conclusion that Brg1 contributes to renal fibrosis by regulating fibroblast phenotype complements, rather than disputes, its roles in other renal cell lineages. Our previously published data argue that deletion of Brg1 in endothelial cells protects the mice from renal injury and fibrosis(22, 48). Additionally, Gong *et al* have shown that Brg1 contributes renal fibrosis by augmenting senescence and suppressing autophagy in tubular cells(16). Combined with our new finding as summarized here, we propose that Brg1 plays cell lineage-specific roles in the kidney to regulate injury and fibrosis. This model seems to be partially supported by our observation that whereas specific elimination of Brg1 from fibroblasts/myofibroblasts attenuates renal fibrosis without altering renal injury, administration of PFI-3, which targets Brg1 non-discriminately, improves both renal injury and fibrosis. Finally, the specificity of PFI-3 towards Brg1 has been questioned previously(71). More recently, a novel and more selective Brg1 inhibitor has been designed and tested(72). It would be of high interest to investigate whether this next-gen Brg1-targeting compound, dubbed IV-255, can efficiently inhibit FMyT and renal fibrosis. These limitations should be carefully considered and tackled so that safe and effective translational strategies can be devised based on the proposed model (Fig.7H).

## Acknowledgements

This work was supported by grants from the National Natural Science Foundation of China (81970638 and 81670618).

## Data Sharing Statement

The data that support the findings of this study are available upon reasonable request. RNA-seq data have been deposited in the PubMed database with the accession number GSE218814, GSE218807, and GSE246869.

## Author contributions

T Zhang, X Wu, and M Kong conceived the project; all authors designed experiments; X Wu, Y Luo, A Kang, J Ni, and M Kong performed experiments, collected data, and analyzed data; all authors wrote and edited the manuscript; T Zhang and X Wu secured funding; M Kong provided supervision and coordination.

## References

1. 2020. Global, regional, and national burden of chronic kidney disease, 1990-2017: a systematic analysis for the Global Burden of Disease Study 2017. Lancet 395:709-733.

2. Chawla, L.S., and Kimmel, P.L. 2012. Acute kidney injury and chronic kidney disease: an integrated clinical syndrome. Kidney Int 82:516–524.

3. Johansen, K.L., Chertow, G.M., Foley, R.N., Gilbertson, D.T., Herzog, C.A., Ishani, A., Israni, A.K., Ku, E., Kurella Tamura, M., Li, S., et al. 2021. US Renal Data System 2020 Annual Data Report: Epidemiology of Kidney Disease in the United States. Am J Kidney Dis 77:A7–A8.

4. Zhou, D., and Liu, Y. 2016. Renal fibrosis in 2015: Understanding the mechanisms of kidney fibrosis. Nat Rev Nephrol 12:68–70.

5. Falke, L.L., Gholizadeh, S., Goldschmeding, R., Kok, R.J., and Nguyen, T.Q. 2015. Diverse origins of the myofibroblast-implications for kidney fibrosis. Nat Rev Nephrol 11:233–244.

6. Mack, M., and Yanagita, M. 2015. Origin of myofibroblasts and cellular events triggering fibrosis. Kidney Int 87:297–307.

7. Quaggin, S.E., and Kapus, A. 2011. Scar wars: mapping the fate of epithelial-mesenchymal-myofibroblast transition. Kidney Int 80:41–50.

8. Zeisberg, E.M., Potenta, S.E., Sugimoto, H., Zeisberg, M., and Kalluri, R. 2008. Fibroblasts in kidney fibrosis emerge via endothelial-to-mesenchymal transition. J Am Soc Nephrol 19:2282–2287.

9. LeBleu, V.S., Taduri, G., O’Connell, J., Teng, Y., Cooke, V.G., Woda, C., Sugimoto, H., and Kalluri, R. 2013. Origin and function of myofibroblasts in kidney fibrosis. Nat Med 19:1047–1053.

10. Kuppe, C., Ibrahim, M.M., Kranz, J., Zhang, X., Ziegler, S., Perales-Paton, J., Jansen, J., Reimer, K.C., Smith, J.R., Dobie, R., et al. 2021. Decoding myofibroblast origins in human kidney fibrosis. Nature 589:281–286.

11. Edeling, M., Ragi, G., Huang, S., Pavenstadt, H., and Susztak, K. 2016. Developmental signalling pathways in renal fibrosis: the roles of Notch, Wnt and Hedgehog. Nat Rev Nephrol 12:426–439.

12. Wu, H., Kirita, Y., Donnelly, E.L., and Humphreys, B.D. 2019. Advantages of Single-Nucleus over Single-Cell RNA Sequencing of Adult Kidney: Rare Cell Types and Novel Cell States Revealed in Fibrosis. J Am Soc Nephrol 30:23–32.

13. Helin, K., and Dhanak, D. 2013. Chromatin proteins and modifications as drug targets. Nature 502:480–488.

14. Hargreaves, D.C., and Crabtree, G.R. 2011. ATP-dependent chromatin remodeling: genetics, genomics and mechanisms. Cell Res 21:396–420.

15. Ross, S., Cheung, E., Petrakis, T.G., Howell, M., Kraus, W.L., and Hill, C.S. 2006. Smads orchestrate specific histone modifications and chromatin remodeling to activate transcription. EMBO J 25:4490–4502.

16. Gong, W., Luo, C., Peng, F., Xiao, J., Zeng, Y., Yin, B., Chen, X., Li, S., He, X., Liu, Y., et al. 2021. Brahma-related gene-1 promotes tubular senescence and renal fibrosis through Wnt/beta-catenin/autophagy axis. Clin Sci (Lond) 135:1873–1895.

17. Lake, B.B., Menon, R., Winfree, S., Hu, Q., Melo Ferreira, R., Kalhor, K., Barwinska, D., Otto, E.A., Ferkowicz, M., Diep, D., et al. 2023. An atlas of healthy and injured cell states and niches in the human kidney. Nature 619:585–594.

18. O’Sullivan, E.D., Mylonas, K.J., Bell, R., Carvalho, C., Baird, D.P., Cairns, C., Gallagher, K.M., Campbell, R., Docherty, M., Laird, A., et al. 2022. Single-cell analysis of senescent epithelia reveals targetable mechanisms promoting fibrosis. JCI Insight 7.

19. Dong, W., Zhu, Y., Zhang, Y., Fan, Z., Zhang, Z., Fan, X., and Xu, Y. 2021. BRG1 Links TLR4 Trans-Activation to LPS-Induced SREBP1a Expression and Liver Injury. Front Cell Dev Biol 9:617073.

20. Khalil, H., Kanisicak, O., Prasad, V., Correll, R.N., Fu, X., Schips, T., Vagnozzi, R.J., Liu, R., Huynh, T., Lee, S.J., et al. 2017. Fibroblast-specific TGF-beta-Smad2/3 signaling underlies cardiac fibrosis. J Clin Invest 127:3770–3783.

21. Kanisicak, O., Khalil, H., Ivey, M.J., Karch, J., Maliken, B.D., Correll, R.N., Brody, M.J., SC, J.L., Aronow, B.J., Tallquist, M.D., et al. 2016. Genetic lineage tracing defines myofibroblast origin and function in the injured heart. Nat Commun 7:12260.

22. Liu, L., Mao, L., Wu, X., Wu, T., Liu, W., Yang, Y., Zhang, T., and Xu, Y. 2019. BRG1 regulates endothelial-derived IL-33 to promote ischemia-reperfusion induced renal injury and fibrosis in mice. Biochim Biophys Acta Mol Basis Dis 1865:2551–2561.

23. Liu, L., Wu, X., Xu, H., Yu, L., Zhang, X., Li, L., Jin, J., Zhang, T., and Xu, Y. 2018. Myocardin-related transcription factor A (MRTF-A) contributes to acute kidney injury by regulating macrophage ROS production. Biochim Biophys Acta Mol Basis Dis 1864:3109–3121.

24. Landegger, L.D., Pan, B., Askew, C., Wassmer, S.J., Gluck, S.D., Galvin, A., Taylor, R., Forge, A., Stankovic, K.M., Holt, J.R., et al. 2017. A synthetic AAV vector enables safe and efficient gene transfer to the mammalian inner ear. Nat Biotechnol 35:280–284.

25. Ikeda, Y., Sun, Z., Ru, X., Vandenberghe, L.H., and Humphreys, B.D. 2018. Efficient Gene Transfer to Kidney Mesenchymal Cells Using a Synthetic Adeno-Associated Viral Vector. J Am Soc Nephrol 29:2287–2297.

26. Piras, B.A., Tian, Y., Xu, Y., Thomas, N.A., O’Connor, D.M., and French, B.A. 2016. Systemic injection of AAV9 carrying a periostin promoter targets gene expression to a myofibroblast-like lineage in mouse hearts after reperfused myocardial infarction. Gene Ther 23:469–478.

27. Nakai, T., Iwamura, Y., and Suzuki, N. 2021. Efficient isolation of interstitial fibroblasts directly from mouse kidneys or indirectly after ex vivo expansion. STAR Protoc 2:100826.

28. Xu, H., Wu, X., Qin, H., Tian, W., Chen, J., Sun, L., Fang, M., and Xu, Y. 2015. Myocardin-Related Transcription Factor A Epigenetically Regulates Renal Fibrosis in Diabetic Nephropathy. J Am Soc Nephrol 26:1648–1660.

29. Hatje, F.A., Wedekind, U., Sachs, W., Loreth, D., Reichelt, J., Demir, F., Kosub, C., Heintz, L., Tomas, N.M., Huber, T.B., et al. 2021. Tripartite Separation of Glomerular Cell Types and Proteomes from Reporter-Free Mice. J Am Soc Nephrol 32:2175–2193.

30. Lv, F., Shao, T., Xue, Y., Miao, X., Guo, Y., Wang, Y., and Xu, Y. 2021. Dual regulation of tank binding kinase 1 (TBK1) by BRG1 in hepatocytes contributes to ROS production. Front Cell Dev Biol 9:745985.

31. Kong, M., Dong, W., Zhu, Y., Fan, Z., Miao, X., Guo, Y., Li, C., Duan, Y., Lu, Y., Li, Z., et al. 2021. Redox-sensitive activation of CCL7 by BRG1 in hepatocytes during liver injury. Redox Biol 46:102079.

32. Kong, M., Dong, W., Xu, H., Fan, Z., Miao, X., Guo, Y., Li, C., Ye, Q., Wang, Y., and Xu, Y. 2021. Choline Kinase Alpha Is a Novel Transcriptional Target of the Brg1 in Hepatocyte: Implication in Liver Regeneration. Front Cell Dev Biol 9:705302.

33. Fan, Z., Kong, M., Miao, X., Guo, Y., Ren, H., Wang, J., Wang, S., Tang, N., Shang, L., Zhu, Z., et al. 2021. An E2F5-TFDP1-BRG1 complex mediates transcriptional activation of MYCN in hepatocytes. Front Cell Dev Biol 9:742319.

34. Liu, L., Zhao, Q., Kong, M., Mao, L., Yang, Y., and Xu, Y. 2022. Myocardin-related transcription factor A regulates integrin beta 2 transcription to promote macrophage infiltration and cardiac hypertrophy in mice. Cardiovasc Res 118:844–858.

35. Dong, W., Kong, M., Liu, H., Xue, Y., Li, Z., Wang, Y., and Xu, Y. 2022. Myocardin-related transcription factor A drives ROS-fueled expansion of hepatic stellate cells by regulating p38-MAPK signalling. Clin Transl Med 12:e688.

36. Shao, T., Xue, Y., and Fang, M. 2021. Epigenetic Repression of Chloride Channel Accessory 2 Transcription in Cardiac Fibroblast: Implication in Cardiac Fibrosis. Front Cell Dev Biol 9:771466.

37. Wu, X., Miao, X., Guo, Y., Shao, T., Tang, S., Lin, Y., Xu, Y., Li, N., and Zhang, T. 2023. Slug enables redox-sensitive trans-activation of LRP1 by COUP-TFII: Implication in antifibrotic intervention in the kidneys. Life Sci:121412.

38. Wu, X., Dong, W., Kong, M., Ren, H., Wang, J., Shang, L., Zhu, Z., Zhu, W., and Shi, X. 2021. Down-regulation of CXXC5 de-represses MYCL1 to promote hepatic stellate cell activation. Front Cell Dev Biol 9:680344.

39. Johnson, B.G., Ren, S., Karaca, G., Gomez, I.G., Fligny, C., Smith, B., Ergun, A., Locke, G., Gao, B., Hayes, S., et al. 2017. Connective Tissue Growth Factor Domain 4 Amplifies Fibrotic Kidney Disease through Activation of LDL Receptor-Related Protein 6. J Am Soc Nephrol 28:1769–1782.

40. Wohlfahrt, T., Rauber, S., Uebe, S., Luber, M., Soare, A., Ekici, A., Weber, S., Matei, A.E., Chen, C.W., Maier, C., et al. 2019. PU.1 controls fibroblast polarization and tissue fibrosis. *Nature* 566:344-349.

41. Lodie, T.A., Reiner, M., Coniglio, S., Viglianti, G., and Fenton, M.J. 1998. Both PU.1 and nuclear factor-kappa B mediate lipopolysaccharide-induced HIV-1 long terminal repeat transcription in macrophages. *J Immunol* 161:268-276.

42. Pongubala, J.M., Van Beveren, C., Nagulapalli, S., Klemsz, M.J., McKercher, S.R., Maki, R.A., and Atchison, M.L. 1993. Effect of PU.1 phosphorylation on interaction with NF-EM5 and transcriptional activation. Science 259:1622-1625.

43. Liang, M.D., Zhang, Y., McDevit, D., Marecki, S., and Nikolajczyk, B.S. 2006. The interleukin-1beta gene is transcribed from a poised promoter architecture in monocytes. J Biol Chem 281:9227–9237.

44. Wang, X., Hu, G., Gao, X., Wang, Y., Zhang, W., Harmon, E.Y., Zhi, X., Xu, Z., Lennartz, M.R., Barroso, M., et al. 2012. The induction of yes-associated protein expression after arterial injury is crucial for smooth muscle phenotypic modulation and neointima formation. Arterioscler Thromb Vasc Biol 32:2662–2669.

45. Zhang, M., Fang, H., Zhou, J., and Herring, B.P. 2007. A novel role of Brg1 in the regulation of SRF/MRTFA-dependent smooth muscle-specific gene expression. J Biol Chem 282:25708–25716.

46. Sun, L., Chen, B., Wu, J., Jiang, C., Fan, Z., Feng, Y., and Xu, Y. 2020. Epigenetic regulation of a disintegrin and metalloproteinase (ADAM) promotes colorectal cancer cell migration and invasion. Front Cell Dev Biol:581692.

47. Bi-Lin, K.W., Seshachalam, P.V., Tuoc, T., Stoykova, A., Ghosh, S., and Singh, M.K. 2021. Critical role of the BAF chromatin remodeling complex during murine neural crest development. PLoS Genet 17:e1009446.

48. Liu, L., Mao, L., Xu, Y., and Wu, X. 2019. Endothelial-specific deletion of Brahma-related gene 1 (BRG1) assuages unilateral ureteral obstruction induced renal injury in mice. Biochem Biophys Res Commun 517:244–252.

49. Zhang, Y., Dees, C., Beyer, C., Lin, N.Y., Distler, A., Zerr, P., Palumbo, K., Susok, L., Kreuter, A., Distler, O., et al. 2015. Inhibition of casein kinase II reduces TGFbeta induced fibroblast activation and ameliorates experimental fibrosis. Ann Rheum Dis 74:936–943.

50. You, E., Jeong, J., Lee, J., Keum, S., Hwang, Y.E., Choi, J.H., and Rhee, S. 2022. Casein kinase 2 promotes the TGF-beta-induced activation of alpha-tubulin acetyltransferase 1 in fibroblasts cultured on a soft matrix. BMB Rep 55:192–197.

51. Huang, J., Chen, Z., Li, J., Chen, Q., Gong, W., Liu, P., and Huang, H. 2017. Protein kinase CK2alpha catalytic subunit ameliorates diabetic renal inflammatory fibrosis via NF-kappaB signaling pathway. Biochem Pharmacol 132:102–117.

52. Soh, U.J., and Low, B.C. 2008. BNIP2 extra long inhibits RhoA and cellular transformation by Lbc RhoGEF via its BCH domain. J Cell Sci 121:1739–1749.

53. Tatsumi, Y., Takano, R., Islam, M.S., Yokochi, T., Itami, M., Nakamura, Y., and Nakagawara, A. 2015. BMCC1, which is an interacting partner of BCL2, attenuates AKT activity, accompanied by apoptosis. Cell Death Dis 6:e1607.

54. Herrmann, D., Straubinger, M., and Hashemolhosseini, S. 2015. Protein kinase CK2 interacts at the neuromuscular synapse with Rapsyn, Rac1, 14–3-3gamma, and Dok-7 proteins and phosphorylates the latter two. *J Biol Chem* 290:22370-22384.

55. Ruzzene, M., Bertacchini, J., Toker, A., and Marmiroli, S. 2017. Cross-talk between the CK2 and AKT signaling pathways in cancer. Adv Biol Regul 64:1–8.

56. Yang, L., Wang, L., Kalfa, T.A., Cancelas, J.A., Shang, X., Pushkaran, S., Mo, J., Williams, D.A., and Zheng, Y. 2007. Cdc42 critically regulates the balance between myelopoiesis and erythropoiesis. Blood 110:3853–3861.

57. Rieske, P., and Pongubala, J.M. 2001. AKT induces transcriptional activity of PU.1 through phosphorylation-mediated modifications within its transactivation domain. *J Biol Chem* 276:8460-8468.

58. Hu, J., Zhang, J.J., Li, L., Wang, S.L., Yang, H.T., Fan, X.W., Zhang, L.M., Hu, G.L., Fu, H.X., Song, W.F., et al. 2021. PU.1 inhibition attenuates atrial fibrosis and atrial fibrillation vulnerability induced by angiotensin-II by reducing TGF-beta1/Smads pathway activation. *J Cell Mol Med* 25:6746-6759.

59. da Silva Antunes, R., Mehta, A.K., Madge, L., Tocker, J., and Croft, M. 2018. TNFSF14 (LIGHT) Exhibits Inflammatory Activities in Lung Fibroblasts Complementary to IL-13 and TGF-beta. Front Immunol 9:576.

60. Yao, L., Shao, W., Chen, Y., Wang, S., and Huang, D. 2022. Suppression of ADAM8 attenuates angiotensin II-induced cardiac fibrosis and endothelial-mesenchymal transition via inhibiting TGF-beta1/Smad2/Smad3 pathways. Exp Anim 71:90–99.

61. Wolk, K., Brembach, T.C., Simaite, D., Bartnik, E., Cucinotta, S., Pokrywka, A., Irmer, M.L., Triebus, J., Witte-Handel, E., Salinas, G., et al. 2021. Activity and components of the granulocyte colony-stimulating factor pathway in hidradenitis suppurativa. Br J Dermatol 185:164–176.

62. Chen, J., Deng, L., Dreymuller, D., Jiang, X., Long, J., Duan, Y., Wang, Y., Luo, M., Lin, F., Mao, L., et al. 2016. A novel peptide ADAM8 inhibitor attenuates bronchial hyperresponsiveness and Th2 cytokine mediated inflammation of murine asthmatic models. Sci Rep 6:30451.

63. Ishizuka, H., Garcia-Palacios, V., Lu, G., Subler, M.A., Zhang, H., Boykin, C.S., Choi, S.J., Zhao, L., Patrene, K., Galson, D.L., et al. 2011. ADAM8 enhances osteoclast precursor fusion and osteoclast formation in vitro and in vivo. J Bone Miner Res 26:169–181.

64. Ding, L., Dolgachev, V., Wu, Z., Liu, T., Nakashima, T., Ullenbruch, M., Lukacs, N.W., Chen, Z., and Phan, S.H. 2013. Essential role of stem cell factor-c-Kit signalling pathway in bleomycin-induced pulmonary fibrosis. J Pathol 230:205–214.

65. Zhang, W., Jia, L., Liu, D.L.X., Chen, L., Wang, Q., Song, K., Nie, S., Ma, J., Chen, X., Xiu, M., et al. 2019. Serum Stem Cell Factor Level Predicts Decline in Kidney Function in Healthy Aging Adults. J Nutr Health Aging 23:813–820.

66. El-Koraie, A.F., Baddour, N.M., Adam, A.G., El Kashef, E.H., and El Nahas, A.M. 2001. Role of stem cell factor and mast cells in the progression of chronic glomerulonephritides. Kidney Int 60:167–172.

67. Gerstenberger, B.S., Trzupek, J.D., Tallant, C., Fedorov, O., Filippakopoulos, P., Brennan, P.E., Fedele, V., Martin, S., Picaud, S., Rogers, C., et al. 2016. Identification of a Chemical Probe for Family VIII Bromodomains through Optimization of a Fragment Hit. J Med Chem 59:4800–4811.

68. Reyes, J.C., Barra, J., Muchardt, C., Camus, A., Babinet, C., and Yaniv, M. 1998. Altered control of cellular proliferation in the absence of mammalian brahma (SNF2alpha). EMBO J 17:6979–6991.

69. Zhou, J., Zhang, M., Fang, H., El-Mounayri, O., Rodenberg, J.M., Imbalzano, A.N., and Herring, B.P. 2009. The SWI/SNF chromatin remodeling complex regulates myocardin-induced smooth muscle-specific gene expression. Arterioscler Thromb Vasc Biol 29:921–928.

70. Stratton, M.S., Haldar, S.M., and McKinsey, T.A. 2017. BRD4 inhibition for the treatment of pathological organ fibrosis. F1000Res 6.

71. Blake, J.F., Burkard, M., Chan, J., Chen, H., Chou, K.J., Diaz, D., Dudley, D.A., Gaudino, J.J., Gould, S.E., Grina, J., et al. 2016. Discovery of (S)-1-(1-(4-Chloro-3-fluorophenyl)-2-hydroxyethyl)-4-(2-((1-methyl-1H-pyrazol-5-yl)amino) pyrimidin-4-yl)pyridin-2(1H)-one (GDC-0994), an Extracellular Signal-Regulated Kinase 1/2 (ERK1/2) Inhibitor in Early Clinical Development. J Med Chem 59:5650–5660.

72. Yang, C., He, Y., Wang, Y., McKinnon, P.J., Shahani, V., Miller, D.D., and Pfeffer, L.M. 2023. Next-generation bromodomain inhibitors of the SWI/SNF complex enhance DNA damage and cell death in glioblastoma. J Cell Mol Med.

